# IL11 stimulates ERK/P90RSK to inhibit LKB1/AMPK and activate mTOR in hepatocytes, the stroma and cancer cells

**DOI:** 10.1101/2022.02.10.479876

**Authors:** Anissa A. Widjaja, Joyce Goh Wei Ting, Sivakumar Viswanathan, Jessie Tan, Shamini G Shekeran, David Carling, Lim Wei Wen, Stuart A. Cook

## Abstract

Interleukin 11 (IL11) stimulates stromal cell activation but also causes hepatocyte metabolic dysfunction. The mechanisms underlying these seemingly unrelated processes are not known. Here we report that IL11-stimulated ERK/P90RSK activity causes the sequential phosphorylation of LKB1 (STK11) at S325 and S428, leading to its inactivation. This leads to a reduction in AMPK activity whilst concomitantly activating mTOR in human fibroblasts, hepatic stellate cells, hepatocytes and cancer cells. In fibroblasts and hepatic stellate cells, IL11-mediated LKB1/AMPK inhibition causes myofibroblast transformation whereas in hepatocytes it inhibits autophagy and fatty acid oxidation and is toxic. Across cell types, the self-amplifying loop of autocrine IL11 activity was inhibited by AMPK activation with metformin, AICAR or 991. In mice on a western diet with fructose, anti-IL11 therapy or hepatocyte-specific deletion of *Il11ra1* rescues LKB1/AMPK activity and reduces NASH. In contrast, restoration of IL11 signalling in hepatocytes of mice with global *Il11ra1* deletion inactivates LKB1/AMPK and exacerbates NASH. These data show that LKB1, an important tumour suppressor and master kinase, is not constitutively active and identify the IL11/LKB1/AMPK/mTOR axis as a point of signalling convergence for epithelial homeostasis, fibrogenesis, immunometabolism and cancer.

## Introduction

Interleukin 11 (IL11) is a little studied cytokine that was recently found to cause ERK-dependent fibroblast-to-myofibroblast transformation and fibrosis across organs (Schafer et al., 2017; Widjaja et al., 2019). Unexpectedly, during our studies of non-alcoholic steatohepatitis (NASH), IL11 was observed to have significant, hepatocyte-dependent metabolic effects (Dong et al., 2021; Widjaja et al., 2019). Genetic or pharmacologic inhibition of IL11 in mice fed a western diet and fructose (WDF) caused generalised improvements in metabolism that included lower body weight, fat mass, liver steatosis and reduced serum triglycerides, cholesterol and glucose levels. Increased hepatic ERK activity was associated with these IL11-induced metabolic changes but the underlying mechanism is unknown (Dong et al., 2021; Widjaja et al., 2019).

We hypothesised that there is a conserved role for IL11-stimulated ERK activity in stromal cells and hepatocytes that underlies both myofibroblast transformation and hepatocyte lipotoxicity / metabolic perturbation. ERK activation in the liver has previously been linked with diet-induced metabolic dysfunction (Jiao et al., 2013; Zheng et al., 2009b) and ERK is a important for activation of both fibroblasts (Du et al., 2008; Schafer et al., 2017) and hepatic stellate cells (HSCs) (Foglia et al., 2019; Widjaja et al., 2019). Intriguingly, IL11 was recently shown to activate rapamycin-sensitive ERK-dependent, mTOR-driven protein translation in fibroblasts (Widjaja et al., 2021a). This nascent discovery raises the question: could the IL11/ERK/mTORC1 axis be a shared mechanism for stromal cell activation and hepatocyte dysfunction and, if so, what is the full signalling pathway?

In this study we investigate whether IL11-stimulated ERK/P90RSK activity can phosphorylate LKB1 (STK11) to inactivate the LKB1/AMPK axis and increase mTOR activity. LKB1 and AMPK are centrally important for both fibroblast activation (Rangarajan et al., 2018; Thakur et al., 2015; Vaahtomeri et al., 2008) and diet-induced liver steatosis (Shaw et al., 2005; Woods et al., 2017). LKB1, a master kinase and tumour suppressor, forms a heterotrimeric complex with MO25 (mouse protein 25) and STRAD (Ste20-related adaptor) and is thought to be constitutively active in the absence of phosphorylation (Zeqiraj et al., 2009). However, inactivation of LKB1 has been reported following dual phosphorylation by ERK/P90RSK in melanoma cells (Zheng et al., 2009a). Here, we use a range of techniques to dissect the effects of IL11 on the ERK/P90RSK/LKB1/AMPK/mTOR axis in fibroblasts, HSCs, hepatocytes, and lung epithelial cancer cells. Furthermore, we examine the effect of genetically or pharmacologically inhibiting IL11 signalling in livers from mice with WDF-induced NASH.

## Results

### An axis of IL11/ERK/P90RSK inhibits LKB1 to activate fibroblasts

We have shown previously that TGFβ1 or IL11 stimulation of human cardiac fibroblasts over a 24 h time course causes disparate p-STAT3 profiles but common biphasic activation of ERK (Widjaja et al., 2021a). We examined whether this pattern was linked with phosphorylation of LKB1 (p-S428; putative inactive LKB1), AMPK (p-T172; active AMPK) or mTOR (S2448; active mTOR) (**Figure 1A and S1**). Following IL11 or TGFβ1 stimulation, LKB1 was phosphorylated at the P90RSK site (S428) in a biphasic manner over the time course. In these experiments, we were not able to assess phosphorylation of endogenous LKB1(S325), the ERK site, as detection was only possible following LKB1 overexpression, as shown below. In stimulated cells, AMPK activity (p-AMPK) was inverse to p-LKB1 levels, with progressive loss in AMPK activity up to 24 h post stimulation. Reciprocal to the decrease in p-AMPK, p-mTOR expression increased over both the IL11 and TGFβ1 time courses.

**Figure 1.**
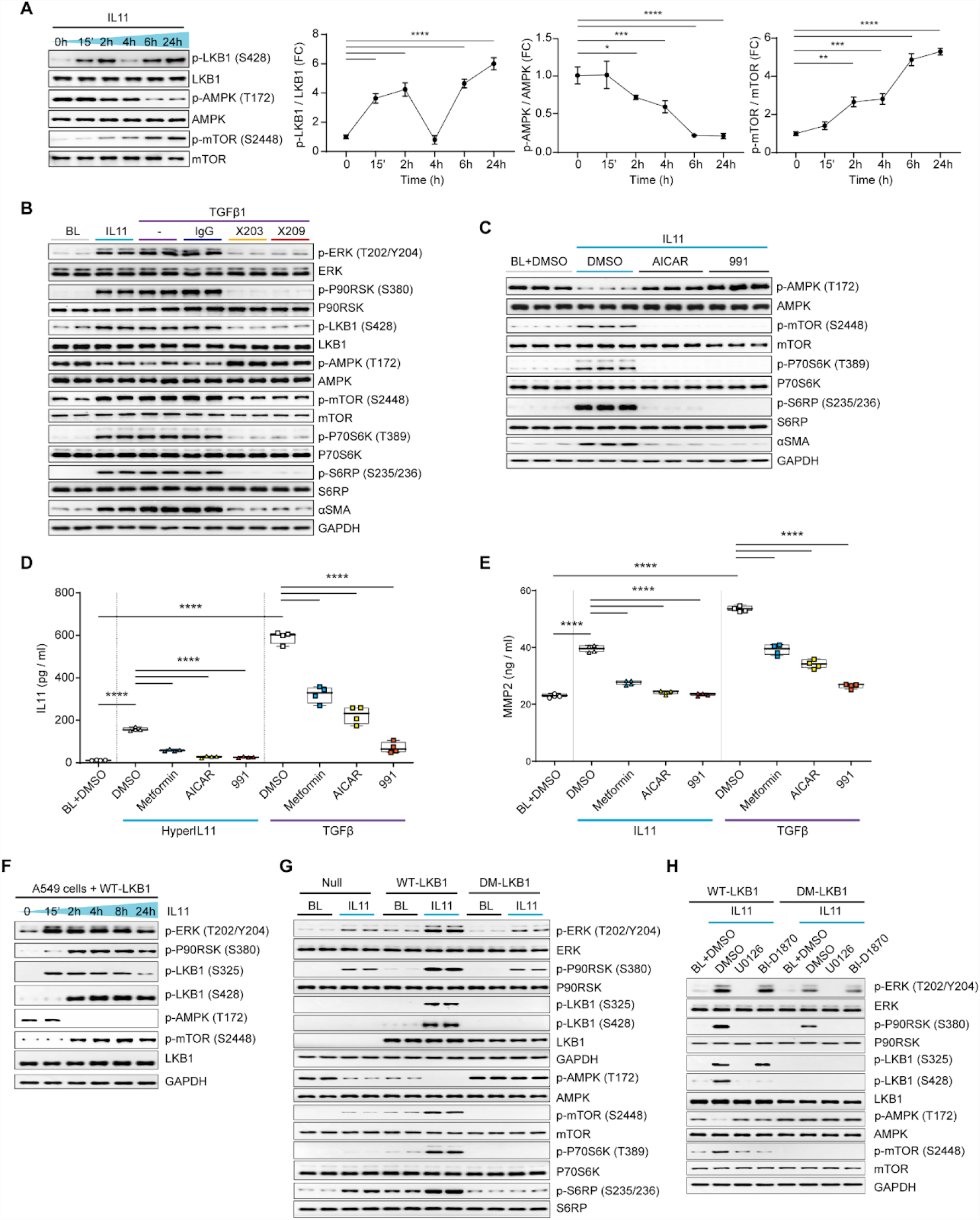
An axis of IL11/ERK/P90RSK inhibits LKB1. (**A**) Western blots (WB) and densitometry analyses of phosphorylated (p-) and total LKB1, AMPK and mTOR in HCFs stimulated with IL11 over a time course; one-way ANOVA with Dunnett’s correction. (**B**) WB of p- and total ERK, P90RSK, LKB1, AMPK, mTOR, P70S6K, S6RP, □SMA and GAPDH levels in HCFs stimulated with IL11 or TGFβ1 alone or in presence of anti-IL11 (X203), antiI-IL11RA (X209) or an IgG control. (**C**) WB of p- and total AMPK, mTOR, P70S6K, S6RP, □SMA and GAPDH levels in IL11-stimulated HCFs in the presence of DMSO, AICAR, or 991. (**D-E**) IL11 (D) and MMP2 (E) concentration in the supernatant of HCFs stimulated with (D) HyperIL11, (E) IL11, or (D-E) TGFβ1 in the presence of DMSO, metformin, AICAR, or 991; one-way ANOVA with Sidak’s correction. (**F**) WB of p-ERK, P90RSK, LKB1, AMPK, mTOR, total LKB1, and GAPDH in A549 cells infected with AAV8-driven expression of wild-type (WT) LKB1 stimulated with IL11 across time (total protein levels **Figure S4**). (**G-H**) WB of p- and total ERK, P90RSK, LKB1, AMPK, mTOR, P70S6K, S6RP, and GAPDH levels from AAV8-driven expression of either null control vector, WT-LKB1 or DM-LKB1-infected A549 cells that are stimulated with IL11 (G) alone or (H) with the addition of DMSO, U0126 or BI-D1870 for 2 hours. (A-H) IL11, TGFβ1, HyperIL11 (10 ng/ml), IgG, X203, X209 (2 μg/ml), AICAR (1 mM), BI-D1870, U0126 (10 μM), DMSO (0.1%), 991 (1 μM), metformin (1 mM); 24-hour stimulation unless otherwise specified. (A) Data are shown as mean±SD (n=3), (D-E) data are shown as box-and-whisker with median (middle line), 25th– 75th percentiles (box) and min-max percentiles (whiskers); *p<0.05, **p<0.01, ***p<0.001,****p<0.0001. BL: Baseline; FC: Fold change

TGFβ1-stimulated ERK activity in fibroblasts is IL11 dependent (Schafer et al., 2017; Widjaja et al., 2021a). We used neutralising antibodies against IL11 (X203) or IL11RA (X209) to assess whether TGFβ1-stimulated p-LKB1 was similarly dependent on IL11 (**Figure 1B**). IL11 and TGFβ1 increased p-ERK, p-P90RSK(S380) and p-LKB1, which was associated with diminished p-AMPK and increased p-mTOR. We extended downstream analyses and, in IL11 or TGFβ1-stimulated cells, we observed levels of p-P70S6K(T389) and its substrate S6RP(S235/236) were increased, as was the mesenchymal marker □SMA (ACTA2).These signalling events were inhibited by X203 or X209, demonstrating the importance of autocrine IL11 activity downstream of TGFβ1, in this context.

MTOR activity is regulated by complex mechanisms and LKB1 controls multiple (>12 (Shackelford and Shaw, 2009)) AMP-related protein kinases. We sought to determine the specific importance of AMPK for mTOR activation by stimulating fibroblasts with IL11 in the presence or absence of the AMPK activating compounds, AICAR or 991 (**Figure 1C**) (Lai et al., 2014). IL11 stimulation (10ng/ml, 24h) caused the expected decrease in p-AMPK and increase in p-mTOR/p-P70S6k/p-S6RP that was associated with higher □SMA expression. In IL11 stimulated cells incubated with either AICAR or 991, p-AMPK levels were maintained, mTOR/P70RSK remained inactive and myofibroblast transformation was inhibited.

We examined if AMPK activity is involved in the self-amplifying loop of IL11 secretion by stimulating fibroblasts with an IL11RA:IL11 fusion protein (HyperIL11 (Schafer et al., 2017)) or TGFβ1 in the presence or absence of AMPK activating compounds (**Figure 1D**). Both HyperIL11 and TGFβ1 induced IL11 secretion (TGFβ1 more so), as expected, and this effect was significantly inhibited by metformin, AICAR or 911. Similarly, IL11-or TGFβ1-induced MMP2 secretion from fibroblasts was reduced by AMPK activation, with a similar profile of inhibition (magnitude of inhibition: 911 > AICAR > metformin) (**Figure 1E**). We have shown previously that IL11 effects in cardiac fibroblasts are critically dependent on ERK (Schafer et al., 2017), which we confirmed for IL11-induced IL11 or MMP2 secretion in the primary fibroblast cultures used here by co-incubation with U0126 (**Figure S2**).

### Phosphorylation of both S325 and S428 in LKB1 is required for its inactivation

To study the relationship between IL11 stimulated p-ERK/p-P90RSK and LKB1 activity we used LKB1 deficient A549 cells for heterologous, AAV8-driven expression of wild type LKB1 (WT-LKB1) or mutant LKB1 (LKB1(S325A and S428A)), which we designated double mutant LKB1 (DM-LKB1). We first confirmed the expression of IL11RA in A549 cells, a lung epithelial cancer cell line (**Figure S3**). We next infected cells with AAV8 encoding WT-LKB1 and stimulated cells with IL11 (10ng/ml, 24 h time course). As seen in primary cells, IL11 increased p-ERK levels in A549 cells, although not biphasically (**Figure 1F**). In these overexpression experiments, the p-LKB1(S325) antibody detected a band co-migrating with LKB1 and, following IL11 stimulation, LKB1 was first (at 15 min) phosphorylated at S325, coincident with increased p-ERK levels, and later (at 2h) phosphorylated at S428, along with P90RSK activation. AMPK activity (p-AMPK) was maintained until both LKB1 sites were phosphorylated at which point p-AMPK levels diminished and p-mTOR levels increased (**Figure 1F and S4**).

A549 cells were then infected with AAV8 encoding WT-LKB1 or DM-LKB1 and stimulated with IL11. Cells expressing WT-LKB1 showed phosphorylation of both S325 and S428 whereas those expressing DM-LKB1 did not, confirming antibody specificity (**Figure 1G**). As compared to control cells, p-ERK and p-P90RSK levels were increased in IL11 stimulated cells expressing WT-LKB1 but to a lesser extent in those expressing DM-LKB1, suggesting a crosstalk effect (Du et al., 2008). In cells expressing WT-LKB1, IL11 stimulation reduced p-AMPK levels and increased p-mTOR/p-P70S6K and p-S6RP levels. In contrast, cells expressing DM-LKB1 maintained high levels of p-AMPK expression after IL11 stimulation and p-mTOR was undetectable.

Our data suggested that, following IL11 stimulation, there is sequential phosphorylation of LKB1 by ERK and then P90RSK (**Figure 1F**) and that dual phosphorylation is needed to fully inhibit LKB1 (**Figure 1F** and **G**). We examined this premise using pharmacologic inhibition of either ERK (U0126) or P90RSK (BI-D1870) in cells stimulated with IL11 (**Figure 1H**). Cells expressing WT-LKB1 showed the expected increases in phosphorylation of ERK/P90RSK/LKB1, decreased p-AMPK, and elevated p-mTOR. In cells expressing DM-LKB1, p-ERK/p-P90RSK was increased following stimulation but the downstream signalling changes were not seen. Cells stimulated with IL11 in the presence of U0126 had no detectable p-ERK/p-P90RSK, no LKB1 phosphorylation, preserved p-AMPK and low p-mTOR (**Figure 1H**). Cells stimulated with IL11 in the presence of BI-D1870 had increased p-ERK but not p-P90RSK, elevated p-LKB1(S325) but not p-LKB1(S428) and preserved p-AMPK and low p-mTOR. These data confirm that LKB1 is phosphorylated sequentially by ERK/P90RSK and that both S325 and S428 need to be phosphorylated to fully inhibit LKB1.

### IL11/ERK activity inhibits LKB1/AMPK causing hepatotoxicity

Pursuing the hypothesis that IL11/ERK is important for LKB1/AMPK activity across cell types we next studied hepatocytes. We first compared the effects of IL6 and IL11 as these cytokines are sometimes portrayed as having overlapping effects in the liver (Schmidt-Arras and Rose-John, 2016). Primary human hepatocytes were stimulated (24 h) with a dose range of IL6 or IL11 which revealed dose-dependent IL6-mediated phosphorylation of STAT3 but not ERK, whereas IL11 dose-dependently increased p-ERK but not p-STAT3 (**Figure 2A**). Across the dose range (1.25-20ng/ml) IL6 had no effect on p-LKB1 or p-AMPK levels. In contrast, IL11 dose-dependently increased p-LKB1 that was accompanied by a reciprocal reduction in p-AMPK.

**Figure 2.**
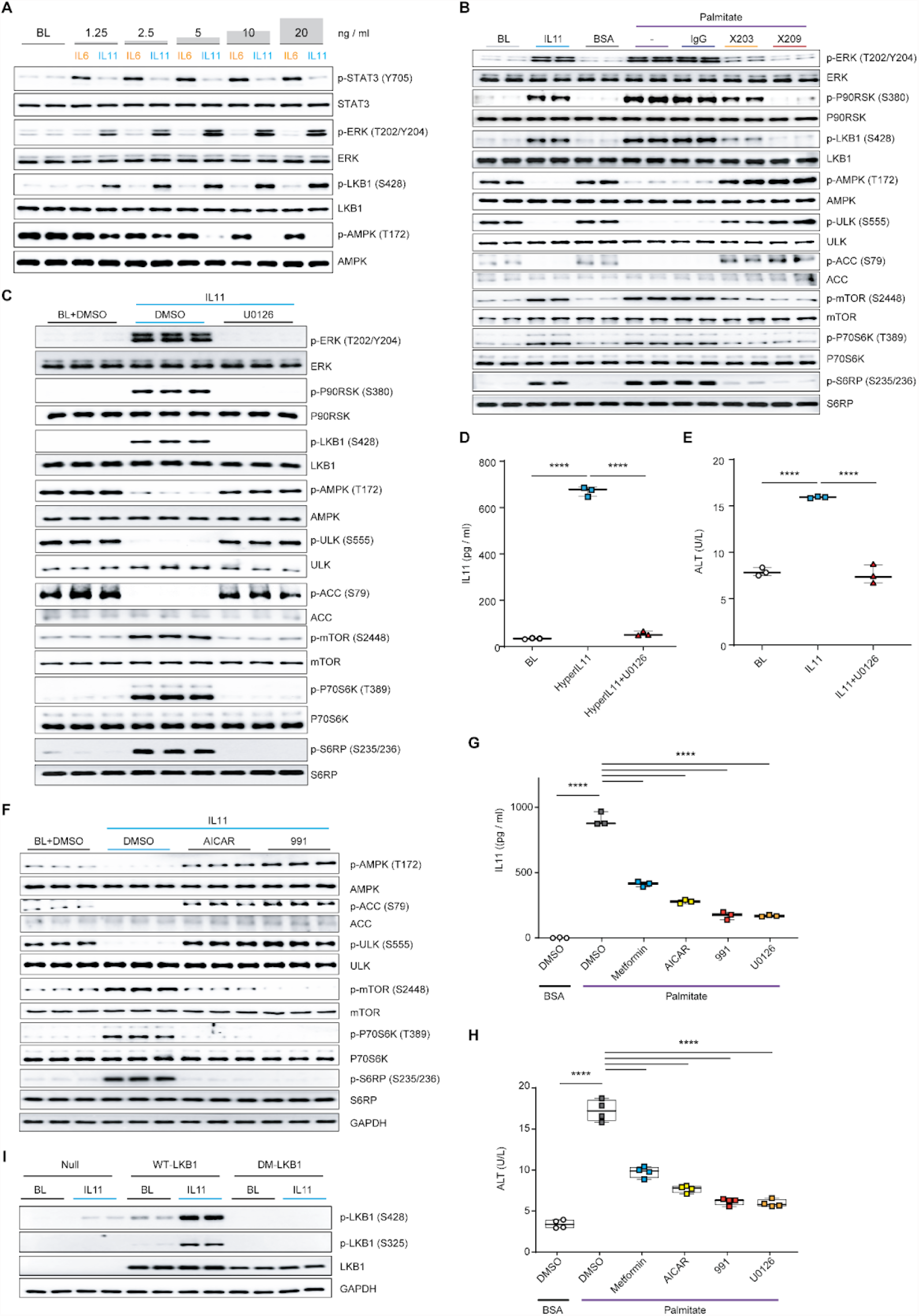
IL11/ERK activity inhibits LKB1/AMPK to cause hepatocytoxicity. (**A**) WB of p- and total STAT3, ERK, LKB1 and AMPK in hepatocytes following a dose range stimulation with either IL6 or IL11. (**B-C**) WB of p- and total ERK, P90RSK, LKB1, AMPK, ULK, ACC, mTOR, P70S6K, and S6RP in hepatocytes stimulated with (B) IL11 or palmitate in the presence of IgG, X203, or X209 or IL11 in the presence of U0126 or DMSO. (**D-E**) Secreted (D) IL11 or (E) ALT in the supernatant of HyperIL11 or (E) IL11-stimulated hepatocytes in the presence of DMSO or U0126; one-way ANOVA with Tukey’s correction. (**F**) WB of p- and total AMPK, ULK, ACC, mTOR, P70S6K, S6RP, and GAPDH in IL11-stimulated hepatocytes in the presence of DMSO, AICAR, or 991. (**G-H**) (G) IL11 and (H) ALT concentrations in the supernatant of palmitate-loaded hepatocytes in the presence of DMSO, metformin, AICAR, 991, or U0126; one-way ANOVA with Dunnett’s correction. (**I**) WB of p- and total LKB1 and GAPDH in AAV8-driven expression of null control vector, WT-LKB1 or DM-LKB1-infected hepatocytes at BL or stimulated with IL11. (A-I) IL11, HyperIL11 (10 ng/ml), IgG, X203, X209 (2 μg/ml), AICAR (1 mM), DMSO (0.1%), 991 (1 μM), metformin (1 mM), U0126 (10 μM); 24-hour stimulation. (D-E, G-H) Data are shown as box-and-whisker with median (middle line), 25th–75th percentiles (box) and min-max percentiles (whiskers) (****p<0.0001). BL: Baseline.

We have described previously an autocrine loop of IL11 activity in steatotic hepatocytes that causes metabolic dysfunction and lipotoxicity (Dong et al., 2021; Widjaja et al., 2019, 2020). To explore this further, we loaded primary human hepatocytes with palmitate in the presence or absence of anti-IL11 (X203) or anti-IL11RA (X209) or stimulated with IL11 (**Figure 2B**). Hepatocytes stimulated with IL11 had increased p-ERK/p-P90RSK/p-LKB1, diminished p-AMPK and increased p-mTOR. Similar to IL11-stimulated hepatocytes, steatotic hepatocytes had lesser LKB1/AMPK activity that was accompanied by decreased p-ACC and p-ULK, consistent with decreased fatty acid oxidation and autophagy (Egan et al., 2011; Ha et al., 1994). Inhibition of IL11 signalling in steatotic hepatocytes with either X203 or X209 abrogated ERK activation, restored p-AMPK levels and increased p-ULK and p-ACC, further confirming the autocrine loop of IL11 activity. The critical and apical role of IL11-stimulated ERK activity for all downstream events was demonstrated using U0126 (**Figure 2C**).

To confirm the existence of autocrine IL11 activity in hepatocytes we stimulated cells with HyperIL11 (**Figure 2D**). HyperIL11 induced IL11 secretion and this effect was inhibited by U0126. IL11 is hepatotoxic (Dong et al., 2021; Widjaja et al., 2021b), which we reaffirmed and also showed to be ERK dependent (**Figure 2E**).

As for fibroblasts (**Figure 1D**), we examined whether restoration of AMPK activity following IL11-induced LKB1 inactivation could rescue downstream events. IL11 stimulation of hepatocytes (10ng/ml, 24h) decreased p-AMPK, p-ACC and p-ULK levels and increased p-mTOR. However, in cells stimulated with IL11 and incubated with either AICAR or 991, p-AMPK, p-ACC and p-ULK levels were maintained and the mTOR/P70RSK/S6RP axis was not activated (**Figure 2F**).

We then determined whether AMPK activity is important for the IL11 feed-forward loop in steatotic hepatocytes (**Figure 2G** and **H**). Metformin, AICAR or 991 all significantly reduced IL11 secretion from lipid laden hepatocytes (**Figure 2G**). Tellingly, treatment of steatotic hepatocytes with these factors also reduced cell death with a similar efficacy profile (metformin < AICAR < 911 (**Figure 2H**). As expected, U0126 strongly inhibited IL11 secretion and cell death in steatotic hepatocytes (**Figure 2G** and **H**). While endogenous LKB1(S325) phosphorylation could not be detected in hepatocyte extracts, we confirmed this event occurred following IL11 stimulation by infecting hepatocytes with WT-LKB1 or DM-LKB1 and then stimulating them with IL11 (**Figure 2I**).

### IL11/ERK activity inhibits LKB1/AMPK to activate hepatic stellate cells

Primary human HSCs were stimulated with a dose range of IL6 or IL11 for 24h (**Figure 3A**). This showed, as for hepatocytes (**Figure 2A**), dose-dependent IL6-mediated phosphorylation of STAT3 but not ERK. In contrast, also as in hepatocytes, IL11 dose-dependently increased p-ERK. Across the dose range (1.25-20ng/ml) IL6 had no effect on p-LKB1 or p-AMPK levels, whereas IL11 dose-dependently phosphorylated LKB1 and inhibited AMPK (**Figure 3A**).

**Figure 3.**
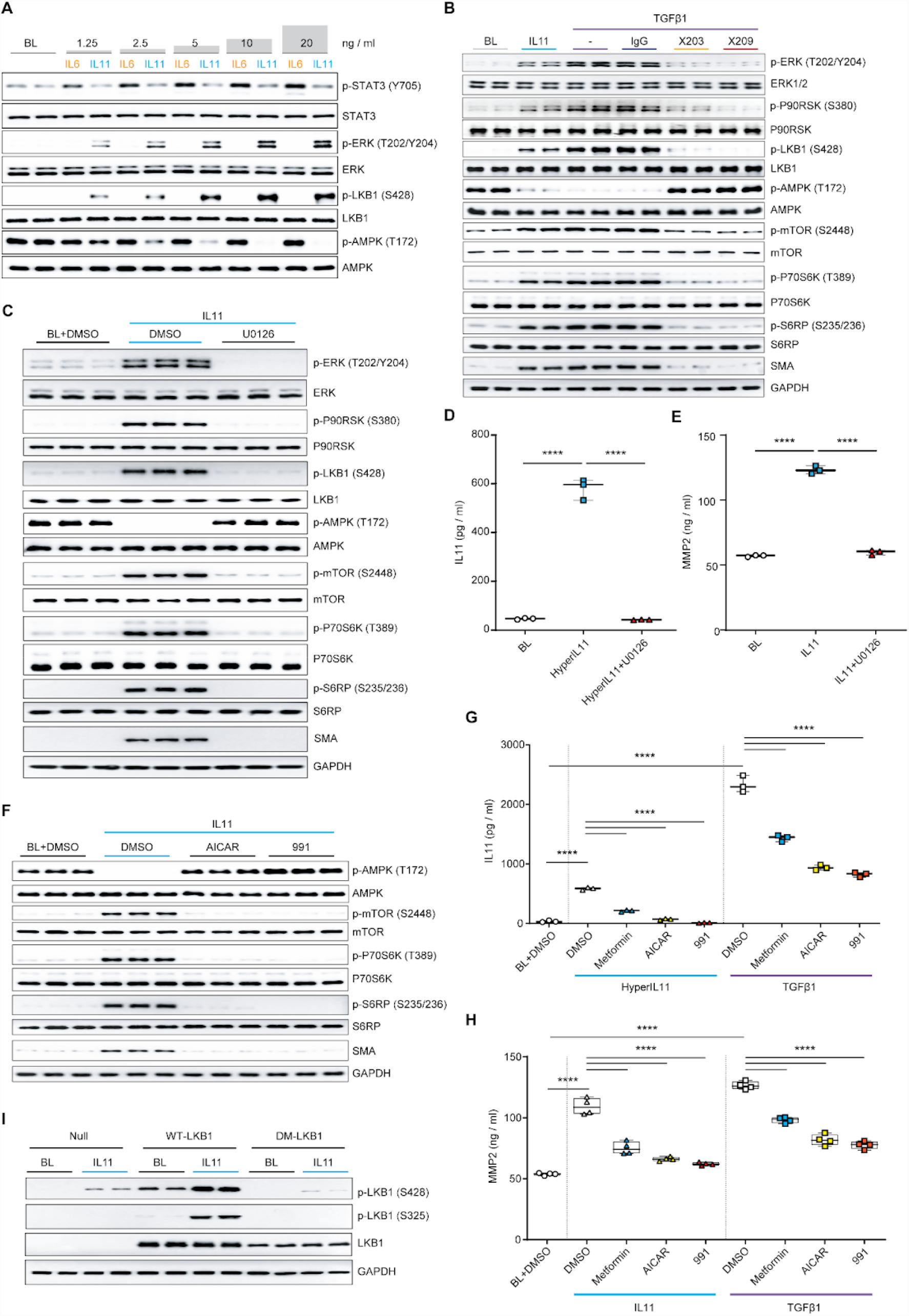
IL11/ERK activity inhibits LKB1/AMPK to activate hepatic stellate cells. (A) WB of p- and total STAT3, ERK, LKB1 and AMPK in HSCs stimulated with increasing doses of either IL6 or IL11. (**B-C**) WB of p- and total ERK, P90RSK, LKB1, AMPK, mTOR, P70S6K, and S6RP in HSCs stimulated with (B) IL11 or TGFβ1 in the presence of IgG, X203, or X209 or (C) IL11 in the presence of U0126 or DMSO. (**D-E**) (D) IL11 or (E) ALT levels in the supernatant from (D) HyperIL11 or (E) IL11-stimulated HSCs in the presence of DMSO or U0126; one-way ANOVA with Tukey’s correction. (**F**) WB of p- and total AMPK, mTOR, P70S6K, S6RP, and GAPDH in IL11-stimulated HSCs with the addition of DMSO, AICAR, or 991. (**G-H**) (G) IL11 and (H) ALT concentrations in the (G) HyperIL11, (H) IL11, or (G-H) TGFβ1-stimulated HSC supernatant in the presence of DMSO, metformin, AICAR, or 991; one-way ANOVA with Sidak’s corrections. (**I**) WB of p- and total LKB1 and GAPDH in AAV8-driven expression of null control vector, WT-LKB1 or DM-LKB1-infected HSCs at BL or stimulated with IL11. (A-I) HyperIL11, IL11, TGFβ1 (10 ng/ml), IgG, X203, X209 (2 μg/ml), AICAR (1 mM), DMSO (0.1%), 991 (1 μM), metformin (1 mM), U0126 (10 μM); 24-hour stimulation. (D-E, G-H) Data are shown as box-and-whisker with median (middle line), 25th–75th percentiles (box) and min-max percentiles (whiskers); ****p<0.0001). BL: Baseline.

Primary human HSCs were incubated TGFβ1 in presence or absence of anti-IL11 (X203) or anti-IL11RA (X209) or stimulated with IL11 (**Figure 3B**). HSCs stimulated with IL11 exhibited increased p-ERK/p-LKB1, diminished p-AMPK and increased p-mTOR/p-P70S6K, p-S6RP. HSCs stimulated with TGFβ1 exhibited similar activation of the ERK/LKB1/AMPK/mTOR axis. Consistent with the requirement for an autocrine loop of IL11 activity downstream of TGFβ1 for HSC activation, anti-IL11 or anti-IL11RA prevented TGFβ1-induced ERK activation, restored AMPK activity and reduced levels of the myofibroblast marker □SMA (**Figure 3B**).

As seen for hepatocytes (**Figure 2C**) the critical and initiating role of IL11-stimulated ERK activity was shown using U0126, which inhibited IL11-mediated ERK activation, all associated downstream signalling and HSC-to-myofibroblast transformation (**Figure 3C**). We stimulated HSCs with HyperIL11 that induced ERK-dependent IL11 secretion, demonstrating the autocrine IL11 loop in HSCs (**Figure 3D**). IL11 induces the secretion of MMP2 from HSCs (Widjaja et al., 2019), and we show here using U0126 that this effect is ERK dependent (**Figure 3E**).

In HSCs, we again assessed the specific role of AMPK downstream of LKB1 in IL11 stimulated cells (**Figure 3F**). Cells stimulated with IL11 and incubated with AMPK-activating compounds had normal p-AMPK levels, did not activate mTOR and did not undergo myofibroblast transformation (**Figure 3F**). Metformin, AICAR or 991 all reduced TGFβ1-or HyperIL11-induced secretion of both IL11 and MMP2 from HSCs with similar profiles of effect (**Figure 3G and H**). Endogenous p-LKB1(325) was not detected in HSCs but we confirmed this event by infecting HSCs with AAV encoding WT-LKB1 or DM-LKB1 in the presence or absence of IL11 stimulation (**Figure 3I**).

### IL11-mediated inhibition of LKB1 contributes to NASH

To translate our findings to the *in vivo* setting we took advantage of samples that we had kept from our previous NASH studies (Dong et al., 2021; Widjaja et al., 2019). We first examined samples from WT mice or mice with conditional deletion/knockout of *Il11ra1* in hepatocytes (CKO) on a western diet with fructose (WDF, 16 weeks) (Dong et al., 2021). We initially probed liver extracts for p-ERK, which was shown before to be elevated in livers of WT mice but not CKO mice on WDF, and substantiated this result, confirming sample integrity (**Figure 4A**).

**Figure 4.**
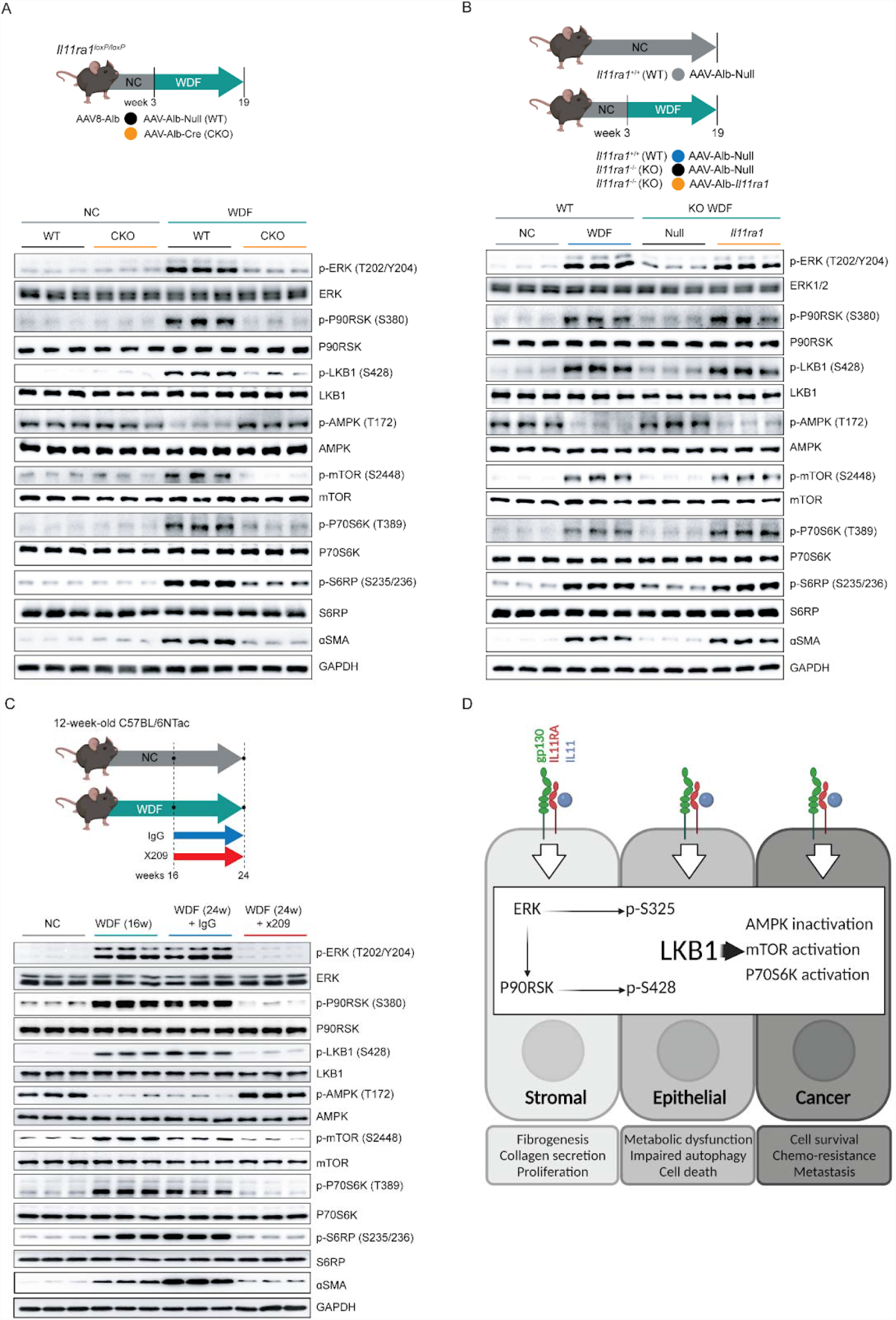
IL11-mediated inhibition of LKB1 contributes to NASH. WB showing hepatic levels of p- and total ERK, P90RSK, LKB1, AMPK, mTOR, P70S6K, S6RP, □SMA and GAPDH from (A) NC or WDF fed-wild-type (WT) or hepatocyte-specific *Il11ra1* knockout mice (CKO), (B) NC or WDF-fed *Il11ra1*^*+/+*^ (WT), WDF-fed *Il11ra1*^*-/-*^ (KO), or WDF-fed KO mice that were previously infected with AAV8-Alb-mbIl11ra1 (full-length membrane-bound *Il11ra1*) in order to specifically restore the expression of *Il11ra1* in hepatocytes, (C) WDF-fed mice that were on therapeutic reversal dosing experiment; mice were treated with 10 mg/kg (2x/week; IP) of either IgG or X209 16 weeks after the start of WDF for a duration of 8 weeks while they were still on continuous WDF feeding. (D) Schematic summarising the role of ERK/P90RSK activity downstream of IL11 stimulation in the phosphorylation of LKB1 at both S325 and S428 sites, leading to its inactivation and the consequent biological effects in stromal, epithelial and cancer cells.

To generate new insights, we then assessed the LKB1/AMPK/mTOR axis and related signalling. This revealed that WT mice on WDF had elevated p-P90RSK/p-LKB1, lesser p-AMPK and increased p-mTOR/p-P70S6K/p-S6RP along with evidence for HSC-to-myofibroblast transformation (□SMA upregulation) (**Figure 4A**). This recapitulates the *in vitro* data in hepatocytes (**Figure 2**), and shows a role for the IL11/ERK/LKB1/AMPK axis in vivo in NASH.

We next studied liver samples from WT mice or mice globally null for *Il11ra1* infected with control AAV8 (WT AAV8) or AAV8 encoding albumin-driven *Il11ra1* expression, which restores IL11RA1 expression in hepatocytes (Dong et al., 2021) (**Figure 4B**). We determined sample integrity by examining p-ERK in liver extracts from WT mice on WDF (16 weeks), which we confirmed was increased. In WT mice on WDF, we again observed elevated p-P90RSK/p-LKB1, lesser p-AMPK and increased p-mTOR and □SMA levels. *Il11ra1* null mice on WDF infected with control AAV8 were protected from NASH and activation of the ERK/LKB1/AMPK axis was not apparent. In contrast, hepatocyte-specific restoration of IL11RA1 expression was associated with increased p-ERK/p-P90RSK/p-LKB1, lesser p-AMPK and these mice developed NASH and metabolic derangement (Dong et al., 2021).

We then studied liver samples from mice with NASH, secondary to WDF for 16 weeks, that were administered a neutralising IL11RA antibody (X209) or an IgG control antibody for a further 8 weeks, while maintained on WDF (Widjaja et al., 2019) (**Figure 4C**). WT mice on WDF for 16 weeks and those receiving IgG by study end (24 weeks) had increased p-ERK, as previously reported, and the expected phosphorylation changes in LKB1/AMPK/mTOR and other proteins. □SMA was upregulated after 16 weeks of WDF and increased further at 24 weeks in mice receiving IgG, showing disease and fibrosis progression. In contrast, the IL11/ERK pathway, and associated downstream changes, was inhibited by administration of X209 from weeks 16 to 24 of WDF and □SMA levels were lower than at 16 weeks, showing disease reversal. We schematically summarise our study findings across cell types in **Figure 4D**.

## Discussion

It is generally accepted that LKB1, an important tumour suppressor and master kinase (Hardie and Alessi, 2013), is constitutively active in the absence of phosphorylation (Shackelford and Shaw, 2009; Zeqiraj et al., 2009). Here we show that IL11-stimulated ERK/P90RSK-mediated LKB1 phosphorylation inactivates LKB1. Dual LKB1 phosphorylation (S325, S428) causing its inactivation has been described before in a B-RAF V600E melanoma cell line (Zheng et al., 2009a). However, other studies found that LKB1 S325 or S428 phosphorylation did not regulate LKB1 activity, perhaps related to the study of individual serine residues (Denison et al., 2009; Sapkota et al., 2002). Our study shows for the first time that LKB1 can be inactivated by IL11-stimulated ERK activity across human primary cell types, as well as in cancer cells.

Some tumour cells, such as A549 (Rapoza et al., 2006), secrete IL11, as do tumour subclones (Janiszewska et al., 2019; Marusyk et al., 2014) and the tumour stroma (Nishina et al., 2021). In Peutz Jeghers syndrome, heterozygous loss-of-function in LKB1 is associated with high levels of IL11 in intestinal polyps and an increased risk of cancers (Ollila et al., 2017). While there is an extensive literature on IL11 in cancer, studies to date have focused on IL11/JAK/STAT signalling (Ernst and Putoczki, 2014; To et al., 2022), which is contentious in some cell types (Widjaja et al., 2021a). Irrespective of this, IL11 has repeatedly been associated with tumour cell proliferation, resistance to chemotherapy and metastatic potential (Ernst and Putoczki, 2014; Janiszewska et al., 2019; Marusyk et al., 2014). We suggest that IL11/ERK-mediated inactivation of LKB1/AMPK, which has been overlooked, is likely important for both tumour cells and the cancer stroma in oncogenesis. Intriguingly, a link between LKB1 mutation, stromal activation and resistance to immunotherapy was recently described (Mazzaschi et al., 2021) and invasive fibroblasts highly express PD-L1 that could act as a decoy (Geng et al., 2019).

In fibroblasts, inhibition of LKB1/AMPK causes myofibroblast transformation and metformin, an indirect AMPK activator, reverses fibrosis (Jiang et al., 2017; Thakur et al., 2015; Vaahtomeri et al., 2008). In hepatocytes, the LKB1/AMPK axis is critical for metabolism, with AMPK inactivation underlying NASH (Long and Zierath, 2006; Shaw et al., 2005; Woods et al., 2017). The dual effect of IL11 in the stroma and epithelium likely explains why its inhibition in NASH reverses not only fibrosis, due to HSC transformation, but also steatosis, due to hepatocyte metabolic dysfunction (Dong et al., 2021; Widjaja et al., 2019). The importance of LKB1/AMPK activity for epithelial cells appears generalisable as renal epithelial cells deficient for LKB1 become steatotic, exhibit mitochondrial dysfunction and deficient LKB1 activity is associated with kidney failure syndromes (Boehlke et al., 2010; Han et al., 2016). It is possible that IL11 activity is important more generally for LKB1-linked pathologies, which requires further study (Han et al., 2016; Hulsurkar et al., 2021; Ollila and Mäkelä, 2011; Sapkota et al., 2002).

IL11 arose in the fish some 700m years ago, is expressed in fish gills and intestine and upregulated in response to viral or bacterial infection (Wang et al., 2005). We speculate that IL11 evolved to protect the epithelial barrier by removing infected epithelial cells and activating scar-forming fibroblasts: parsimonious ERK-mediated inactivation of LKB1/AMPK underlying both these effects. This ancient defence mechanism appears inappropriately and chronically activated in some human diseases (e.g. pulmonary fibrosis), in states of overnutrition and in cancer. We wonder if some of the beneficial effects of metformin on fibrosis (Rangarajan et al., 2018), and perhaps other phenotypes, might be related to inhibition of autocrine IL11 activity. To conclude, we identify IL11 as a key immunometabolic factor that controls LKB1/AMPK and mTORC1 activity across multiple human cell types to drive diverse cellular pathologies important for human diseases.

## Material and Methods

### Ethics statements

All experimental protocols involving human subjects (commercial primary human cell lines) have been performed in accordance with the *ICH Guidelines for Good Clinical Practice*. As written in their respective datasheets, ethical approvals have been obtained by the relevant parties and all participants gave written informed consent to ScienCell from which primary human cardiac fibroblasts, primary human hepatocytes, and primary human hepatic stellate cells were commercially sourced.

Animal studies were carried out in compliance with the recommendations in the *Guidelines on the Care and Use of Animals for Scientific Purposes* of the *National Advisory Committee for Laboratory Animal Research* (NACLAR). All experimental procedures were approved (SHS/2014/0925 and SHS/2019/1482) and conducted in accordance with the SingHealth Institutional Animal Care and Use Committee.

### Antibodies

Phospho ACC Ser79 (11818, CST), ACC (3676, CST), phospho AMPK Thr172 (2535, CST), AMPK (5832, CST) E-Cadherin (3195, CST), phospho-ERK1/2 Thr202/Tyr204 (4370, CST), ERK1/2 (4695, CST), GAPDH (2118, CST), IgG (11E10, Aldevron), neutralizing anti-IL11 (X203, Aldevron), neutralizing anti-IL11RA (X209, Aldevron), IL11RA (ab125015, abcam, IF), phospho LKB1 Ser325 (gifted by Prof. Dario Alessi, University of Dundee), phospho LKB1 Ser428 (3482, CST), LKB1 (3047S), phospho-mTOR Ser2448 (2971, CST), mTOR (2972,CST), phospho-p70S6K Thr389 (9234, CST), p70S6K (2708, CST), phospho P90RSK Ser380 (11989, CST), RSK (9355, CST), phospho-S6 ribosomal protein Ser235/236 (4858, CST), S6 ribosomal protein (2217, CST), □SMA (19245, CST), SNAI1 (3879, CST), phospho-STAT3 Tyr705 (4113, CST), STAT3 (4904, CST), phospho ULK1 Ser555 (5869, CST), ULK1 (8054, CST), mouse HRP (7076, CST), rabbit HRP (7074, CST).

### Recombinant proteins

Commercial recombinant proteins: human IL6 (206-IL, R&D Systems), human insulin (I9278, Sigma), human TGFβ1 (PHP143B, Bio-Rad).

Custom recombinant proteins: Recombinant human IL11 (rhIL11, UniProtKB:P20809) that was synthesised without the signal peptide. HyperIL-11 (IL11RA:IL-11 fusion protein) was constructed using a fragment of IL11RA (amino acid residues 1–317; UniProtKB: Q14626) and IL-11 (amino acid residues 22–199, UniProtKB: P20809) with a 20 amino acid linker (GPAGQSGGGGGSGGGSGGGSV) (Dams-Kozlowska et al., 2012). All custom recombinant proteins were synthesised by GenScript using a mammalian expression system.

### Chemicals

AICAR (A9978, Sigma), BI-D1870 (S2843, Selleck Chemicals), DMSO (D2650, Sigma), ex229 (compound 991, S8654, Selleck Chemicals), Glucose (G7021, Sigma), Metformin (PHR1084, Sigma), Palmitate (P5585, Sigma), U0126 (9903, CST).

### Cell culture

All cells were grown and maintained at 37°C and 5% CO_2_. The culture of primary human cardiac fibroblasts (HCFs), primary human hepatic stellate cells (HSCs), and primary human hepatocytes were described previously (Widjaja et al., 2019, 2021a). Human lung epithelial carcinoma cell line (A549; CCL-185, ATCC) were grown and maintained in DMEM complete medium which contains 10% FBS (10500064, Gibco) and 1% P/S (15140122, Gibco). The growth medium was renewed every 2–3 days and cells were passaged at 80% confluence, using standard trypsinization techniques. All experiments with primary cells were carried out at low cell passage (<P3). Cells were serum-starved overnight in basal media prior to stimulation. Cells were stimulated with different treatment conditions and durations, as outlined in the main text or figure legends. Stimulated cells were compared to unstimulated cells that have been grown for the same duration under the same conditions, but without the stimuli. AICAR, BI-D1870, 991, and metformin were dissolved in 100% DMSO and added to the media in 1:1000 dilution to get a final DMSO concentration of 0.1%. Palmitate was conjugated in fatty acids free BSA in the ratio of 6:1; 0.5% BSA was used as control.

### MOI/transfection exp in A549

A549 cells were seeded in 6-well plates at a density of 2 × 10^5^ cells/well. To determine the optimal multiplicity of infection (MOI), A549 cells were first infected with Ad-CMV-NULL (1300, Vector Biolabs), AAV-WT-LKB (Ad-STK11, 1563, Vector Biolabs) and AAV-DM LKB (Ad-hLKB S325A/S428, ADV-224574, Vector Biolabs) at an MOI of 5, 10 or 20 and incubated for 2 or 24 hours at 37°C. After infection, cells were washed with PBS and the efficiency of gene expression was determined by immunoblotting for LKB1, based on this analysis an MOI of 20 chosen for all experiments.

### Animal models

Animal models were first described in detail in (Widjaja et al., 2019) and (Dong et al., 2021). Mice were housed in temperatures of 21-24□ with 40-70% humidity on a 12 h light/12 h dark cycle and provided with food and water ad libitum.

### Mouse models of NASH

8-week-old C57BL/6N, *Il11ra1*^*-/-*^ mice, and *Il11ra1*^*loxP/loxP*^ and their respective control were fed western diet (D12079B, Research Diets) supplemented with 15% weight/volume fructose in drinking water (WDF) for 16-24 weeks as outlined in the main text/figure legends. Control mice received normal chow (NC) and tap water.

*Il11ra-floxed (Il11ra1*^*loxP/loxP*^) *mice*

6-8-week-old male homozygous *Il11ra1*-floxed mice (Dong et al., 2021) were intravenously injected with 4×10^11^ genome copies (gc) of AAV8-Alb-Cre virus (CKO) or a similar amount of AAV8-Alb-Null virus. Both controls and CKO mice were fed WDF or NC three weeks following virus administration. *Il11ra1-*deleted mice (KO)

6-8-week-old male *Il11ra1*^*-/-*^ mice (Schafer et al., 2017) were intravenously injected with 4×10^11^ gc of AAV8-*Alb*-*Il11ra1* virus to induce hepatocyte specific expression of mouse *Il11ra1*. As controls, both *Il11ra1*^*-/-*^ mice and their wildtype littermates (*Il11ra1*^*+/+*^) were intravenously injected with 4×10^11^ gc *AAV8-Alb-Null* virus. 3 weeks after virus injection, mice were fed with WDF or NC.

*In vivo administration of anti-IL-11 or anti-IL11RA monoclonal antibodies*

Mice were fed with WDF for 16 weeks to induce NASH and then treated with either 10 mg/kg (2X/week via intraperitoneal injection) of anti-IL11RA (X209) or IgG isotype control (11E10) for 8 weeks while they were on continuous WDF feeding.

### Colorimetric assays and enzyme-linked immunosorbent assay (ELISA)

Alanine Aminotransferase (ALT) activity in the cell culture supernatant was measured using ALT Activity Assay Kit (ab105134, Abcam). The levels of IL11 in equal volumes of cell culture media were quantified using Human IL11 Quantikine ELISA kit (D1100; R&D Systems). Both assays were performed according to the manufacturer’s protocol.

### Statistical analysis

All statistical analyses were performed using GraphPad Prism software (version 9). One-way ANOVA with Dunnett’s correction was used when several conditions were compared to one condition), one-way ANOVA with Tukey’s correction was used when several conditions were compared to each other. One-way ANOVA with Sidak’s correction was used when different stimulants and corresponding inhibitors were conducted concurrently and compared against one common baseline condition. The criterion for statistical significance was set at P□<□0.05.

## ACKNOWLEDGEMENTS

We thank Prof. Lewis Cantley for discussion relating to this manuscript and Prof. Dario Alessi for kindly supplying the p-LKB1(S325) antibody. This research is supported by the National Medical Research Council (NMRC), Singapore STaR awards (NMRC/STaR/0029/2017), NMRC Centre Grant to the NHCS, MOH□CIRG18nov□0002, Tanoto Foundation. A.A.W. is supported by NMRC/OFYIRG/0053/2017. The authors would like to acknowledge the technical support of J. Dong and B.L.George.

## AUTHOR CONTRIBUTIONS

S.A.C and A.A.W. conceived and designed the study. A.A.W., J.G.W.T., S.V., and J.T. performed *in vitro* cell culture, cell biology and molecular biology experiments. J.G.W.T. and S.G.S. performed *in vivo* studies. A.A.W., D.C., L.W.W., and S.A.C. analysed the data. A.A.W., L.W.W., and S.A.C. prepared the manuscript with input from co-authors.

## COMPETING INTERESTS

A.A.W. and S.A.C. are co-inventors of the patent applications: US Patent App. 16/865,259 (Treatment and prevention of metabolic diseases) and US Patent App. 16/748,698 (Treatment of hepatotoxicity). S.A.C is a co-inventor of the patent applications: WO/2017/103108 (TREATMENT OF FIBROSIS), WO/2018/109174 (IL11 ANTIBODIES), WO/2018/109170 (IL11RA ANTIBODIES) and a co-founder and shareholder of Enleofen Bio PTE LTD, a company that developed anti-IL11 therapeutics, which were acquired for further development by Boehringer Ingelheim. The remaining authors declare that the research was conducted in the absence of any commercial or financial relationships that could be construed as a potential conflict of interest.

## Supplementary Figures

**Figure S1.**
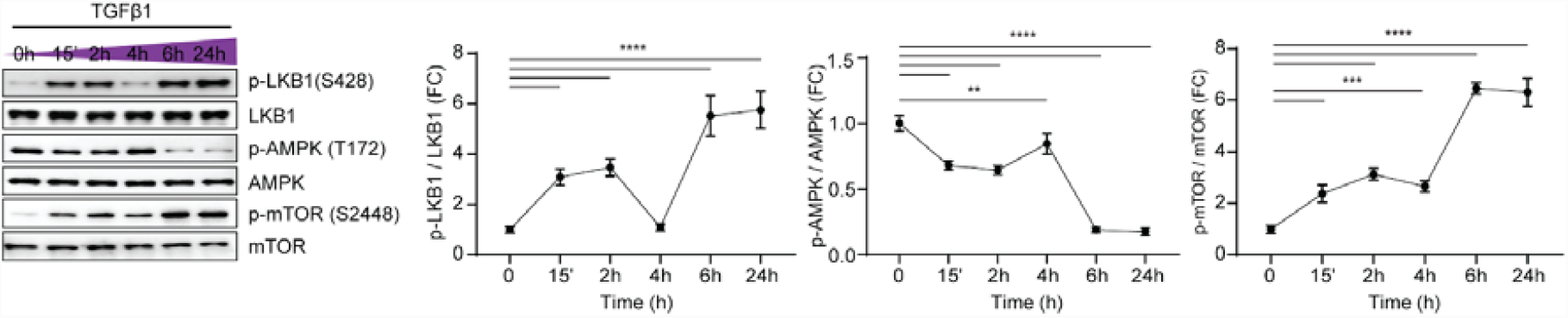
TGFβ1 stimulation inactivates LKB1 in cardiac fibroblasts. Western blot and densitometry analyses of phosphorylated (p-) and total LKB1, AMPK and mTOR in human cardiac fibroblasts stimulated with TGFβ1 (10 ng/ml) over a time course. Data are shown as mean±SD (n=3); **p<0.01, ***p<0.001, ****p<0.0001. FC: Fold change

**Figure S2.**
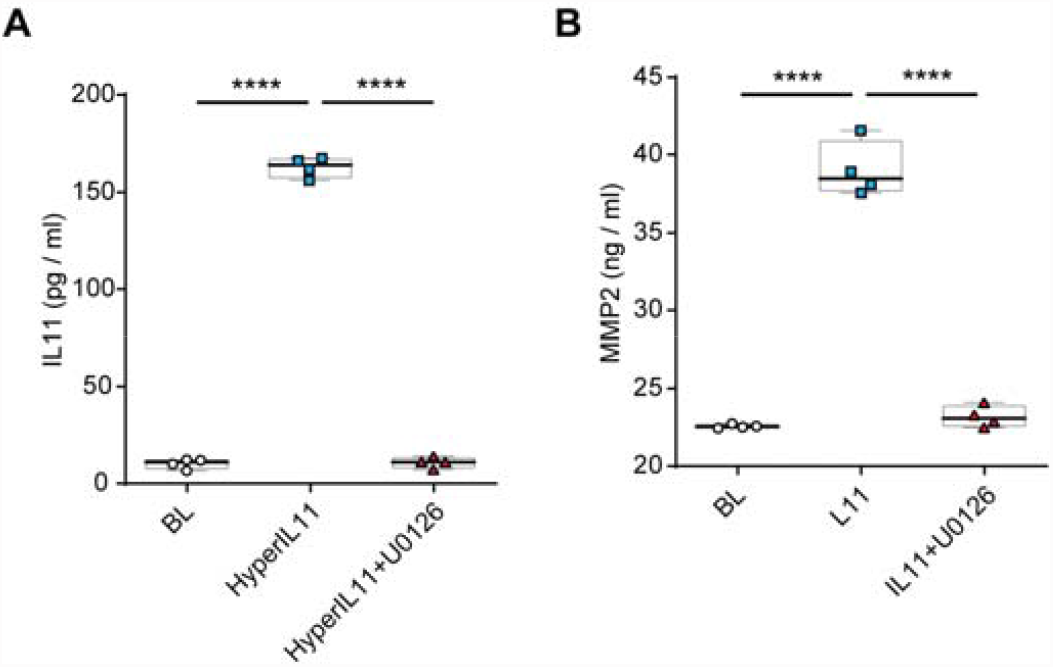
Secretion of IL11 and MMP2 from cardiac fibroblasts is dependent on ERK activity. (**A-B**) Secreted (A) IL11 or (B) MMP2 from human cardiac fibroblasts stimulated with (A) HyperIL11 or (B) IL11 in the absence or presence of U0126. All conditions were carried out in the presence of 0.1% DMSO. Data are shown as box-and-whisker with median (middle line), 25th–75th percentiles (box) and min-max percentiles (whiskers); one-way ANOVA with Tukey’s correction ****p<0.0001. BL: Baseline

**Figure S3.**
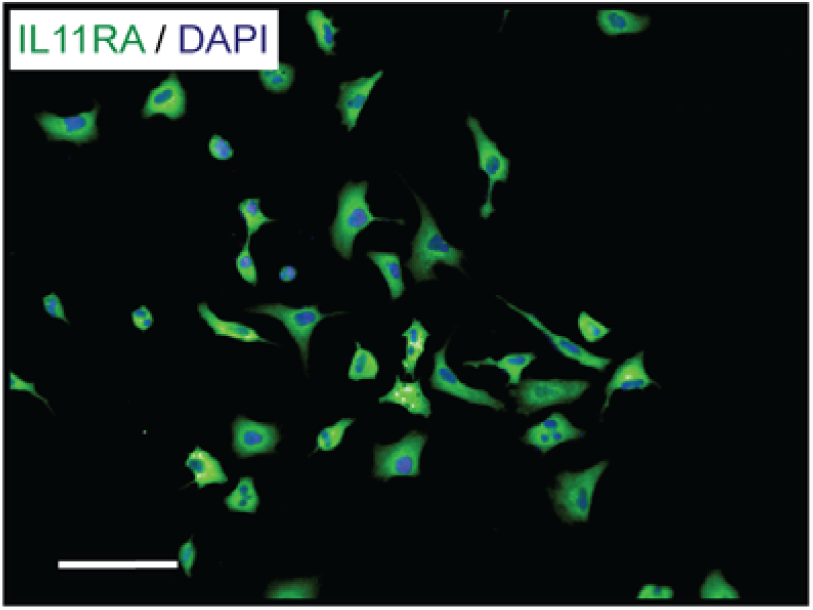
A549 cells express IL11RA. Immunofluorescence image of IL11RA expression in A549 cells at baseline (representative dataset from n=3/group, scale bar, 100 μm)

**Figure S4.**
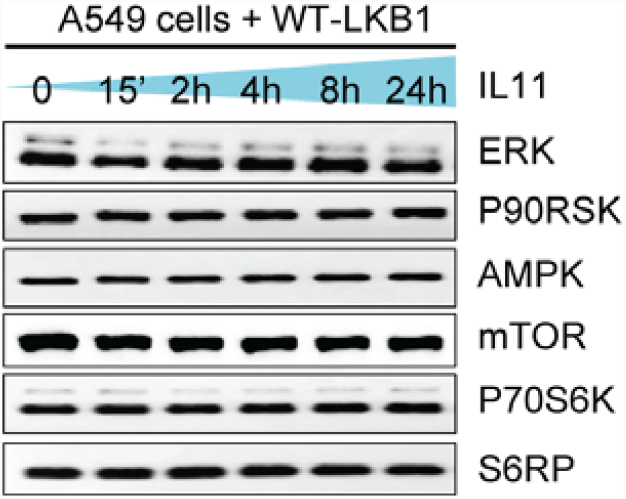
Sequential phosphorylation of LKB1 by ERK and P90RSK. WB of total ERK, P90RSK, AMPK, mTOR, P70S6K and S6RP in IL11(10ng/ml)-stimulated A549 cells infected with AAV8-driven expression of wild type (WT)-LKB1 across time. Experiments were conducted concurrently with **Figure 1F**.

